# Dust or disease? Perceptions of influenza in rural Southern Malawi

**DOI:** 10.1101/470856

**Authors:** Mackwellings Phiri, Kate Gooding, Ingrid Peterson, Ivan Mambule, Spencer Nundwe, Meredith McMorrow, Nicola Desmond

## Abstract

**Background:** Influenza virus infections cause between 291 243 and 645 832 deaths annually, with the highest burden in low-income settings. Research in high-income countries has examined public understanding of influenza, but there is little information on views and behaviours about influenza in low-income countries. We explored communities’ ideas about the severity, causes, prevention and treatment of influenza in Chikwawa district, Malawi.

**Methods:** We conducted 64 in-depth interviews with parents of children aged <5 years, and 7 focus groups with community health workers, parents, and traditional healers. Data were analysed thematically and using a framework matrix to compare views between groups.

**Results:** Respondents held varied ideas about influenza, and many were uncertain about its causes and treatment. Some parents, traditional healers and health workers thought influenza was not severe because they felt it did not cause death or limit activities, but others disagreed. Many saw influenza as a symptom of other conditions, especially malaria and pneumonia, rather than as a disease of its own. Most mentioned dust as the main cause of influenza and believed influenza could be prevented by cleaning the home thoroughly. Treatment seeking for influenza followed different stages, usually starting with home remedies followed by purchasing drugs from groceries and then visiting a health centre. Seeking a clinician tended to be triggered by severe symptoms like high fever or difficulty breathing, and suspicions of malaria or pneumonia. Community health workers provide health education for communities, but some lacked understanding of influenza.

**Conclusion:** Our findings suggest uncertainty about the causes and control of influenza among parents and varied levels of understanding among health providers. Strengthening the capacity of community health workers to provide relevant information about influenza prevention and treatment could address parents’ interest in further information and support informed health seeking and engagement with future influenza interventions.

## Background

Influenza is a global health concern, causing between 291 243 and 645 832 deaths annually (1). Low-income countries bear the highest burden (2–5). Studies in Africa indicate high levels of influenza-associated hospitalization (2,3,6). A high prevalence of co-infections and poor living conditions contribute to poor outcomes from influenza infections in low-income countries (7–9). Given this burden, there has been increased attention on control strategies, including vaccination (3,8) and promotion of personal hygiene [10].

Understanding public views about influenza is crucial for developing effective control strategies (10). The ways that people perceive a disease affects acceptance and utilization of control measures (7,11), so knowledge of community attitudes can support effective design of health programmes (11,12). There is considerable research on public perceptions of influenza in high-income countries (10,13,14), and growing research in middle-income countries. For example, research in South Africa, Kenya and India found varied understandings of influenza, including knowledge of viruses and contagion but also explanations that differed from biomedical models, such as influenza being caused by dust, weather and dirt (15–17). Little is known about perceptions and behaviours about influenza in low-income countries (18).

This paper examines community perceptions and behaviour about influenza in Malawi, a low-income country in Southern Africa. Preliminary research suggests knowledge and beliefs about influenza vary considerably in Malawi (19), but there is little detailed information on the range of beliefs about the causes of influenza or methods for prevention and treatment. In Malawi, influenza control strategies are not currently included within the Health Sector Strategic Plan (20). In common with most African countries, patients are not routinely tested for influenza (21), and antiviral treatment is not available (22). During the 2009 pandemic, vaccination was offered to pregnant women, children and health workers in some districts (19,23), and research indicates a potential role for vaccination of pregnant women (24) and people with HIV (8). However further research is needed to identify effective and feasible influenza control options (8,19,24). Potential introduction of an influenza vaccination also requires understanding of acceptability, and there is limited evidence on influenza vaccine acceptability within the Malawian context (25). To support effective design and introduction of control strategies, we examined community perceptions and health seeking about influenza and views on influenza vaccination.

## Methods

### Study Setting

This study took place in Chikwawa, a rural district in Southern Malawi with an under-five mortality rate of 66 per 1000 live births [25] and a poverty rate of 82% [26]. Chikwawa has a high burden of malaria, schistosomiasis and malnutrition [27–29]. Less than 10% of men and 5% of women aged 15-49 have completed primary school, over 20% of men and 45% of women are illiterate [30], and most households are involved in small-scale farming (32,33) with few involved in paid labour (34).

Our study was conducted in parallel to a community-based Phase IV trial examining the impact of malaria exposure on immune response to influenza vaccination in children aged 6-59 months [26].

### Data collection

We conducted in-depth interviews with parents of study-eligible children in 41 households. We purposefully sampled parents of study-eligible children based on consent or refusal to enrol their children in the vaccine trial, and from villages with varied health centre access. We also conducted a further 20 in-depth interviews using critical incident narratives (CIN) with parents who had brought a child aged less than five years to a health centre with either severe acute respiratory infection or influenza-like illness. CIN interviewees were purposefully selected to include those whose children had different severe acute respiratory infections and influenza like illnesses, and attending different levels of health centre (village, primary and secondary).

Seven focus group discussions (FGDs), each with 9-12 participants, were held: four with parents of children aged less than five years (split between men and women, and between parents from more socially included and excluded households), one with traditional healers, and two with health workers. We defined socially included households as those participating in any social group in the community, for example, money lending or church groups that are typical institutions in the Malawi setting (36). Socially excluded referred to those not involved in such groups. The health workers were all community health workers (Health Surveillance Assistants, or HSAs), except for one more senior officer working on immunization.

All interviews and FGDs included discussion about understanding of influenza: perceived severity, symptoms, causes, treatment and overlap with other conditions. In interviews, we asked about general approaches to management of influenza, and about specific recent individual episodes of illnesses that participants attributed to influenza. CIN asked parents to recount details of a recent illness episode, from initial recognition of symptoms to arriving at a health centre (37).

Data were collected by two authors (MP and SN) with support from a social scientist (KG). Debriefing meetings were held soon after every data collection episode to review emerging findings and identify areas needing further investigation. Interviews lasted 26-105 minutes (with most being 45-60 minutes), and FGDs averaged 90 minutes (ranging from 89-116 minutes). All interviews and FGDs were conducted in the local language (Chichewa) and audio recorded.

### Data analysis

Audio recordings were transcribed verbatim, translated into English, and imported into NVIVO10 software (QSR, Melbourne, Australia) to facilitate organization and analysis. Transcripts were read and re-read for familiarization with the data. Transcripts were coded inductively. KG, MP and SN each coded an initial sample of transcripts, before comparing interpretations. We then generated an integrated coding frame that was used to code further transcripts, iteratively adapting the frame as new data were analysed. A framework matrix was developed to compare views among parents, and between parents, community health workers and traditional healers.

Every participant provided written informed consent. Ethical approval for the study was obtained from the Malawi College of Medicine (16-004) and Liverpool School of Tropical Medicine (P.11/15/1828) research ethics committees. The U.S. Centers for Disease Control and Prevention relied on the local review by the Malawi College of Medicine.

## Results

In examining perceptions of influenza and associated health seeking, we identified themes related to symptoms, severity, causes, prevention and treatment. In discussing influenza with participants, we used the Chichewa word ‘chimfine’, which is the usual translation of influenza in Chichewa (38). However, the same term ‘chimfine’ is also used for the common cold, sneezing, and for a runny nose and hay fever (38). This broad definition needs to be considered in interpreting the results below, and is an issue we return to in the discussion. When quoting participants, we used the word flu because that is the term a lay person would normally use in English.

### Understanding of influenza as a disease and its symptoms

Parents, community health workers and traditional healers all commonly described influenza as a condition that makes people sneeze and unable to breathe normally.

> *Flu is a disease that makes your nose run frequently. The disease also makes you unable to breathe properly.* [Interview, female parent]
>
> *Flu is a disease that makes you sneeze, and as you sneeze, it makes you have a runny nose*. [FGD, health worker]
>
> *Flu is a disease that comes with regular sneezing, causes the nose to be stuffy, and causes you to have a runny nose*. [FGD, traditional healer]

Participants also distinguished influenza symptoms from those of conditions such as malaria, pneumonia or cough (malungo, chibayo or chifuwa, respectively, in Chichewa), citing nasal congestion, sneezing and a runny nose as more associated with influenza.

> *Flu causes the nose to be stuffy and closed, whereas pneumonia causes fast breathing and in-drawing of the chest, and we tell malaria through fever, vomiting and diarrhoea*. [Interview, female parent]
>
> *When you have flu, it makes you have a regular runny nose or sneeze frequently. But if you have cough you only cough but you can’t have a runny nose*. [FGD, health worker]
>
> *[Flu and pneumonia] are different. Flu causes your nose to be stuffy whereas pneumonia causes pain in your ribs when you breathe*. [FGD, traditional healer]

However, other participants identified overlaps in symptoms between influenza and other conditions. For example, some parents and community health workers saw fever and body pains as common to both influenza and malaria, and fever and difficulty breathing as typically associated with influenza and pneumonia.

> *Malaria is another disease with the same symptoms as flu; you feel body pains as well as fever*. [FGD, female parent]
>
> *Flu and pneumonia have similar symptoms, because when you have pneumonia, you gasp for air, and you also gasp for air when you have flu*. [Interview, female parent]
>
> *When you have flu, you experience fever just as when you are sick with malaria; and when you have malaria, you feel body pains just as when you have flu.* [FGD, health worker]
>
> *Most children admitted to hospital because of pneumonia, when they come to the hospital, they have signs like those of flu: high fever and gasping*. [FGD, senior health worker]

Among those parents who described overlapping symptoms for influenza and other illnesses, some identified influenza and other conditions as separate diseases but thought only a doctor could distinguish them.

> *Both malaria and flu come with fever, but you can’t tell [on your own] whether you have malaria or flu unless you go to a hospital and are tested by a doctor. Then you would know that this is malaria or flu*. [Interview, female parent]

However, for other parents this overlap in symptoms suggested that influenza was an early indicator of other conditions such as malaria and pneumonia, rather than a disease in its own right.

> *Flu and malaria are not really different because they go together. You start with flu, then eventually, after some body pain and headache, you realize that the flu you had was actually malaria*. [Interview, female parent]

Those who saw influenza as an indicator of other conditions said they usually had influenza before diagnosis with malaria, leading to the perceived association:

> *Malaria and flu go together because when a child is sick with malaria is also sick with flu. The child is already showing symptoms of flu before malaria becomes visible; the body feels hot*. [Interview, female parent]

Similarly, some community health workers believed influenza was a symptom of malaria because *“most children with flu frequently also test positive for malaria.”* Some health workers also associated influenza with measles, and a small number of traditional healers also described influenza as a symptom of other conditions

> *…flu sometimes can be a sign of another infection like measles…sometimes it can be a primary infection, but usually it comes as a sign…* [FGD, health worker]
>
> *Flu is the origin of different diseases. You first have flu before you have measles…that shows flu is a sign alerting us that we are going to be sick with some disease: measles…and… malaria*. [FGD, traditional healer]

However, while some parents described these understandings of influenza as a disease and how it is recognized, others indicated doubts about what influenza really was.

> *What the word ‘chimfine’ (flu) means? It’s hard for me to tell what flu is…we only get sick with flu…but we don’t know what it is*. [Interview, female parent]
>
> *We only know the word ‘chimfine’ (flu)…but we don’t understand what it is*. [Interview, female parent]

### Types and names for influenza

Parents described several types of influenza. Common flu (termed Chimfine cha wamba) was the kind that parents said normally made them sick. Several parents also mentioned bird flu (Chimfine cha mbalame) and swine flu (Chimfine cha nkhumba), but said they had only heard about these conditions and not experienced them.

> *We only hear people discuss bird flu… we see posters at the district hospital showing people infected with bird flu, which show bird flu is more dangerous than the flu we know*. [Interview, female parent]
>
> *I hear about bird flu, swine flu, but I only know of the flu that we usually get sick with.* [Interview, male parent]

Some parents also described malaria-like flu (“Chimfine cha malungo”), which referred to a form of influenza that resembled malaria in symptoms, and that could develop into malaria.

> *Malaria-like flu starts with sneezing, then you have headache, and high fever follows. Then we know that you are going to have malaria*. [Interview, female parent]

### Perceived severity of influenza

Perspectives on severity of influenza varied between respondents. Some parents felt influenza was not as dangerous as other conditions, such as malaria, measles and pneumonia. This was primarily because parents felt influenza did not seriously affect daily activities.

> *Flu is better compared to other diseases, because you can work when you have flu, but can you work when you have malaria? You can’t. That makes flu better than other diseases.* [Interview, female parent]

Some traditional healers shared this view that influenza was not serious because it had less impact on activities than other illnesses:

> *We think of flu as a mild condition because, even though you are sneezing, you can go out and work. You take treatment while you are working unlike with other conditions where you lie down when you are sick.* [FGD, traditional healer]

Alongside the minimal impacts on activities, some parents felt influenza was less serious than other illnesses because they perceived it as not causing death. Death was only considered a risk through other conditions related to influenza.

> *It has never happened and I have never heard that someone died because of flu; if someone with flu dies, maybe it’s because of other conditions that are associated with it*. [FGD, male parent]

However, other parents felt influenza was just as serious as other conditions, significantly affecting activities or bringing a risk of death. In relation to impact on activities, some said influenza affected work either directly, or through caring responsibilities for others who are sick.

*You can’t do chores if someone has flu because you must look after them. If you returned from work and found that the condition of the child has worsened, you stop going to work; you take the child to the hospital. That means some of the things you need to do won’t be completed because of the sick person*. [Interview, male parent]

In relation to health risks, difficulty breathing due to nasal congestion was the main danger that parents associated with influenza.

> *The danger of flu is that it affects breathing. If you are infected with flu, you can’t breathe normally, your breathing is insufficient.* [Interview, male parent]

While parents had different views about severity, nearly all health workers said influenza was dangerous, particularly because of its impact on breathing and potential to progress to other conditions.

> *You can’t breathe properly when you have flu, and failure to breathe properly may lead to other illnesses; you may end up having pneumonia or malaria.* [FGD, health worker]

Although most health workers thought potential to cause other conditions made influenza dangerous, others saw influenza as only dangerous in combination with other conditions.

> *Unless combined with the other illnesses, flu alone does not cause death. Yes, flu is also a problem, but its danger doesn’t amount to one that can cause death.* [FGD, health worker]

Participants saw severity of influenza as varying between groups. Most parents and health workers thought influenza was particularly dangerous for infants because difficulty breathing due to influenza limited sucking and eating. Health workers also mentioned a risk that breathing problems could cause death for young children.

> *Flu is very dangerous for children because shortness of breath due to stuffy nose makes sucking hard for the child. The child also doesn’t want to eat because of breathing problems*. [Interview, female parent]
>
> *Flu…is a serious problem especially for children because it makes them gasp because air passages are blocked…because the nostrils are closed, the child has problems sucking milk…* [FGD, health worker]
>
> *Flu is serious for small children because it makes them gasp for air, and if a child gasps for air, the child can easily lose consciousness, and repeated loss of consciousness is a serious complication for a child because the child can die.* [FGD, health worker]

Older people, pregnant women and people with pre-existing health problems were also seen as at particular risk from influenza. Parents and health workers said older people were vulnerable to influenza because their natural defence against infection was impaired.

> *Maybe ‘soldiers’ [immunity] in their body are weak, that’s why they are often prone to catching flu*. [FGD, female parent]
>
> *The group at most risk of flu is the elderly. If an older person was infected with flu, they would suffer a lot because their body protection against diseases is weakened.* [FGD, health worker]

Health workers stated that people who already had serious conditions like cancer would find it harder to recover from influenza, and so would pregnant women:

> *Expectant mothers are already weak; if they catch flu, they’ll become even weaker. Also, those who are suffering from critical conditions like diabetes and cancer; if they caught flu, their state would get even worse.* [FGD, health worker]

### Perceived causes of influenza

Participants had several ideas about causes of influenza, and the same individuals often mentioned multiple causes. The most commonly identified cause was dust, often reflecting the experience of sneezing after inhaling dust:

> *Flu infection comes about because of dust. If you are sweeping a mud floor without applying water, and if you breathe in the rising dust, you may get flu.* [Interview, female parent]
>
> *I think flu is caused by dust because when you are sweeping in the house and the dust enters your noses, you start sneezing as though you sniffed pepper powder.* [FGD, traditional healer].

Linked partly with the idea of dust, people also thought influenza was caused by lack of hygiene, which they defined as not cleaning the house or not washing clothes or beddings.

*Poor hygiene can make you catch flu. If a child lies where there is dust, or if you don’t wash a child’s clothes, or if the child sleeps under a dirty bed net, the child will definitely catch flu because the things the child uses are dirty.* [Interview, female parent]

While most participants saw dust as causing influenza directly, others linked the role of dust to germs. They used the Chichewa word ‘kachirombo’, which covers viruses but also bacteria and parasites (hence our use of the broad term ‘germs’ in translation). A few parents thought germs responsible for influenza were in the dust, a view shared by one traditional healer and health worker.

> *There must be germs that cause flu, germs that we can’t see with our naked eyes. Why are we saying germs? Because if it was just dust, we wouldn’t be taking medication. We take medication to fight the germs.* [FGD, traditional healer]
>
> *What I know as a health worker is that dust is the main cause of flu. There are germs in the dust that cause flu*. [FGD, health worker]

Another cause identified by some participants was the idea of influenza as spread from other people, often expressed in relation to proximity and congestion.

> *If you have flu and share a room with other people, you can pass it on to them, and they pass it on to others, and eventually you all become infected*. [Interview, female parent]
>
> *Others say [you catch flu] because of sleeping in a congested area, where you breathe in the air that others breathe out, and that’s a risk*. [FGD, traditional healer]

Transmission between people was only explicitly linked to spread of germs by the more senior health worker (who used the English term virus), and one parent and traditional healer.

> *We know flu as one of the infections that are caused by viruses, and flu spreads from one person to another*. FGD, senior health worker]
>
> *…let us just say flu is a disease caused by a germ that spreads when we stay close to the person that is infected*. [Interview, female parent]
>
> *…what happens is that…if you have flu and share a bed with others, as you sneeze, you pass it on to another person and all of you catch it, because it is a germ that spreads from one person to another*. [FGD, traditional healer]

A further cause of influenza identified by many participants was the weather. However, there were different ideas about what kind of weather caused influenza, and about how different kinds of weather led to influenza. Many people associated influenza with winter (May – July in Malawi), when the weather is cold and dry, with typical temperatures of around 4 to 10 degrees Celsius (39).

> *When the cold season is on, there are so many cases of flu; you hear, one person after the other, “I have flu,” and you have others sneezing. Flu is very common during the cold periods we mentioned – May, June and July*. [FGD, female parent]

This association with cold weather was partly because of an idea that being cold could in itself contribute to influenza.

> *I also think that cold weather can make a child have flu*. [Interview, female parent]
>
> *Here in the villages flu affects us because some of us do not have proper clothing such as jumpers for protection from cold, unlike in summer when we are not affected by cold*. [FGD, female parent]

However, the link between cold weather and influenza also related to the earlier idea of dust causing influenza because dry and windy conditions during the cold season meant increased levels of dust:

> *This time around, if you look around, you will find that most children have flu. This is because of the wind that’s blowing. In the rainy season, there’s no dust because the ground is wet, and there are fewer cases of flu*. [FGD, health worker]

Cold weather was also linked to influenza through the idea of influenza as spread between people. One traditional healer suggested that cold weather made people lie closer together, fuelling the spread of influenza.

> *When it is cold everyone wants to sleep where it’s warm. If you are sharing a bed with other people, you want to sleep close to one another. If you have influenza and are sneezing, as you want to sleep close to each other, the one next to you breathes in the air from your sneezing, and the flu spreads in the house*. [FGD, traditional healer]

In contrast, others linked influenza to hot weather and saw the heat of the sun as leading to influenza.

> *…[the child caught flu] because of so much sun because children like playing in the sun; when they are playing in the sun, their body takes in so much heat, and the body then starts to feel hot*. [Interview, female parent]

Beyond these immediate causes, some people pointed to wider contextual factors that contributed to influenza, for example housing conditions were seen as increasing the risk of person-to-person contagion.

> *[Flu is common in the villages] also because the houses we are living in do not have windows, not enough air goes in. So we can catch flu easily because the houses do not have windows and the air circulating in the house is not good*. [FGD, health worker]

Although participants described these causes of influenza, as with the understanding about what influenza is, many parents indicated uncertainty about what causes influenza.

> *[What causes influenza?] Maybe I should have asked the same question when you asked if I had any questions…I don’t know what causes flu*. [Interview, female parent]
>
> *I’m not sure what causes flu. I only discover that I’m having flu, but I’m not sure what causes it*. [Interview, female parent]

### Strategies for preventing influenza

Methods of preventing influenza often followed perceived causes. For example, those who identified dust as a cause tended to mention steps to prevent exposure to dust, such as dampening earth floors before sweeping, regular maintenance of floors, not working in dusty conditions and keeping children away from dust. These strategies were mentioned by both parents and health workers.

> *When someone catches flu in the house, we ask ourselves: is it because we are not cleaning the house? Or because we are not sealing the floor? So, we do all these things to reduce the risk of the disease.* [Interview, female parent]
>
> *We advise them that if the house is full of dust, when sweeping they should first sprinkle some water so the dust doesn’t rise. We also tell them to clean rooms that are not frequently used*. [FGD, health worker]

As well as cleaning to remove dust, most people saw cleaning and hygiene as important for preventing influenza more generally, and this included laundry and bathing.

> *You can protect your child from flu by bathing them or washing their clothes frequently… that way, the child can’t be infected with flu because you are hygienic*. [Interview, female parent]

Linked with the idea of cold weather as leading to influenza, some parents felt wearing warm clothes could prevent influenza.

> *A child like this one needs proper clothing against cold weather for protection from flu. You need to clothe the child with a sweater, a hat and socks, so that the child feels warm*. [Interview, female parent]

Regarding the idea of influenza as spread between people, a few parents mentioned keeping away from people infected with influenza, and one mentioned washing hands after nose-blowing.

> *To ensure the person with flu doesn’t spread it to others, you should not stay close to them. The person should also stand where the wind blows rather than where it comes from, so that you are not in contact with the air that the person breathes out*. [FGD, male parent]
>
> *When you have flu, to stop spreading it to others, you should wash your hands after blowing your nose, because you might interact with friends after blowing your nose without washing hands and spread it*. [FGD, male parent]

Similarly, one health worker discussed advising people to cover their mouths when sneezing as a person-to-person transmission control measure.

> *…some adults don’t cover their mouths when they have flu and are sneezing, so we encourage them to have a piece of cloth with them…* [FGD, health worker]

In line with ideas about influenza as a symptom of other conditions, some health workers felt influenza could be prevented indirectly by preventing those other conditions, for example, through vaccination against pneumonia.

> *Sometimes flu signals another disease, especially pneumonia, and there are vaccines against pneumonia. Pentavalent vaccine protects children from virus-caused pneumonia and PCV protects children from bacteria-caused pneumonia. Children are unlikely to get flu if they are protected from such infections*. [FGD, senior health worker]

While some parents identified ways to prevent influenza, others felt incapable of preventing the disease. For some, this related to a feeling that transmission could not be stopped:

> *…preventing flu is hard because flu is like an air-bone disease; the moment one in the house contracts it, then you know that all of you in the house are going to be infected. So it’s difficult to say that this is how we can prevent it. We are failing to prevent it*. [FGD, male parent]

For others, the feeling that influenza could not be prevented stemmed from uncertainty about its causes and a perceived lack of information:

> *I don’t know how to prevent flu because I don’t know how I contract it. Maybe if you the doctors tell us what to do then we will know how to prevent it.* [Interview, female parent]

### Approaches for managing and treating influenza

Participants reported several approaches for managing and treating influenza.

Some parents visited a health centre, especially when influenza symptoms were severe, and often because of a suspicion that influenza may have progressed to other conditions perceived as more serious. This approach was supported by health workers. If influenza presented with high fever for instance, suspected malaria was often the reason for going to hospital. Respondents used the Chichewa term ‘chipatala’, which is translated as hospital but can also refer to primary health centres.

> *You don’t go to hospital instantly when you have flu…when it invites those other conditions, headache or pneumonia, that’s when you go to hospital*. [FGD, male parent]
>
> *If a child has flu and it is affecting breathing to a point that the child gasps for air, or the child has fever too and can’t eat, it’s a critical sign that the child should no longer be managed at home…because usually there is pneumonia involved if it gets to that point.* [FGD, health worker]

For situations when influenza was less severe, parents described using home remedies such as bathing in warm water, holding a warm cloth on the nose, taking a salt solution, and frequent fluid intake.

> *If you drink water frequently or take a warm bath or squeeze your nose with a warm cloth or add some salt to the water that you drink, you can treat flu in no time.* [Interview, female parent]

Health workers and traditional healers also mentioned drinking water.

> *…when they come to the hospital…we tell them that if they have flu they should go and take water frequently*. [FGD, health worker]
>
> *Taking water frequently is another treatment for flu because the water cleans body organs and makes them relaxed*. [FGD, traditional healer]

There were different ideas about how drinking water could help. One set of ideas related to having enough water in the body, including the idea of cleaning organs mentioned by this healer above, and other ideas related to water dissolving things or through effects on the blood. Other ideas related to respiratory symptoms, including reducing itching or dryness in the throat and opening the nose.

> *When you have flu and drink plenty of water, the water dissolves everything in the body, and the body functions properly if it has enough water*. [FGD, heath worker]
>
> *Taking water reduces flu because the blood runs properly [because] you have sufficient water in the body*. [FGD, traditional healer]
>
> *Water prevents your throat from drying. I have been infected with flu before. Sometimes you can’t breathe properly because the nose is blocked. But when you take water the nose clears and you are able to breathe normally*. [FGD, male parent]

Another home remedy, discussed by a small number of parents and some traditional healers, was using herbs to help manage influenza by clearing the nose.

> *Others collect leaves that have a very strong scent. They press them and sniff the scent*. [FGD, male parents]
>
> *If they have flu, we give them some herbs to sniff. After sniffing the herb, you find that they’re feeling better*. [FGD, traditional healer]

Some participants mentioned going to traditional healers, especially when something originally presumed to be influenza persisted and so they suspected witchcraft instead.

> *When you visit the hospital and the condition does not improve, you see herbalists because they may have put the child under a spell*. [Interview, female parent]
>
> *When I visited, they gave him medication. After a week, he started vomiting and having diarrhoea. I was so confused and decided to go to a traditional healer. If you have been to the hospital and it’s not helping, you visit traditional healers. The way the condition was, I thought it required seeing a traditional healer*. [Interview, female parent]

Traditional healers also reported people visiting them about influenza, usually after having gone to the hospital.

> *A person needs to go to the hospital first. If they have been to the hospital and the condition doesn’t improve, that’s when we help them*. [FGD, traditional healer]

However, most parents said they did not visit traditional healers, and perceived them as unreliable and causing conflicts in households.

> *I felt that if I went there [traditional healer], it might create enmity… between me and my family…because healers might tell you that your mother is responsible for the sickness of your child*. [Interview, female parent]
>
> *Traditional healers do not provide help, they only make people hate each other. They are full of lies. They might tell you that you are a witch bewitching my child when it’s not like that*. [Interview, female parent]

As well as home remedies, many parents mentioned purchasing drugs from the grocery, such as paracetamol and antibiotics. Health workers also mentioned that parents often took antibiotics before seeing a clinician.

> *When my children had flu, I gave them paracetamol to relieve the headache and fever they were having. I bought the paracetamol from a grocery shop.* [Interview, female parent]
>
> *To manage the problem of flu…you take pills, Bactrim…we buy it from the grocery or we go to the hospital, so that the body may get some relief. [Interview, female parent]*
>
> *… some … believe in certain drugs, when you ask them, “What drugs have you taken?” you hear, “I bought some injectable drugs and pills.” If you ask what these injectable drugs or pills were, you find they are talking about Penicillin or Amoxicillin. They go to shops and ask for these drugs*. [FGD, health worker]

As the first parent’s comment suggests, paracetamol was seen as relieving the effects of influenza, rather than providing a cure. This view was linked to an idea that influenza has no treatment, described in relation to both drug and non-drug options.

> *There is no treatment for flu; you take paracetamol because sneezing or blowing the nose causes headache, so you take the drug just for relief from the headache.* [Interview, female parent]
>
> *What we use as treatment for flu, like ‘luni’ (Gynandropsis Gynandra), we use them just to get some relief*. [Interview, female parent]
>
> *[Bathing in hot water and applying a hot cloth] help but not really. You feel relief only for a short while. The next day the problem starts again, because you are not taking treatment for the disease, because the treatment is not there*. [FGD, female parent]

Previous advice from health centres or clinicians contributed to this idea that influenza has no treatment, and encouraged home management. Parents and some health workers described clinicians advising patients that there was no treatment, and only prescribing paracetamol.

> *When you visit the hospital, they tell you that flu has no treatment, they only give you drugs for relieving the pain you feel like a headache.* [Interview, female parent]
>
> *…when they come to the hospital…if they have flu and are also feeling headache, we give them paracetamol to stop the headache*. [FGD, health worker]

Only the more senior health worker explicitly linked the lack of treatment to the idea of influenza as caused by a virus.

> *We know that flu is one of the diseases that are caused by viruses, and for any disease caused by a virus there is no treatment*. [FGD, senior health worker]

Related to the idea of influenza as having no treatment, people talked about influenza as self-limiting, saying “it ends on its own”. This contributed to some parents saying they just wait when they have influenza, rather than looking for any medication or treatment.

> *It’s better to just stay at home, because with flu, after seven days you find that you are OK. [Interview, female parent]*

The uncertainty seen in relation to causes and prevention of influenza was also found in relation to treatment. Some parents and health workers described not knowing how to treat influenza.

> *…we get sick with flu but we don’t know how it’s treated, and we wonder whether flu is also a disease*. [FGD, female parent]
>
> *Flu is hard to treat because we are not sure what its treatment is*. [FGD, health worker]

### Sources of information for influenza

Most parents mentioned friends and relatives, especially elder parents, as the usual source of information about influenza, including information on home-based management.

> *I heard from friends, my husband and my husband’s brother; they were the ones who said I should cover the child with a cold cloth*. [Interview, female parent]
>
> *Our parents are the ones who teach us about using herbs. They tell us there were no hospitals in the past; they were relying on herbs*. [FGD, male parent]

Parents mentioned health workers as another source of information on health matters, both through hospital visits and public health information sessions in the community. However, most parents felt health workers prioritized information about other diseases and did not disseminate information about influenza.

> *I have never received any information about flu whenever I have been to the hospital, maybe information about cholera, malaria and other diseases…but I have never had information about flu from the hospital*. [Interview, female parent]
>
> *There’s no information [about flu] that’s given because when you tell them [health worker] you have flu, they just write in your card and prescribe medication, but they don’t give you any information as to what you should do*. [Interview, female parent]

This lack of information may have contributed to the uncertainty about influenza indicated in previous sections.

### Views on an influenza vaccine

Parents had not heard about an influenza vaccine before learning about it through the influenza vaccine trial or our meetings with them. Having learnt about it through the trial, most parents were very positive about the vaccine and described vaccination as providing protection and as the most effective means of stopping influenza.

> *The most effective way is a vaccine, because if you give a child a vaccine, the child will be safe from the disease [influenza] regardless of whether the child plays in dirt or the house is dusty*. [Interview, female parent]
>
> *The vaccine they are trying out will be good because if you get it, it means you are protected in advance.* [Interview, female parent]
>
> *We would love if the government would ensure that this vaccine reaches out to everyone, so we can stop this disease*. [FGD, female parent]

Health workers were confident that communities would welcome an influenza vaccine. However, they felt there should be adequate sensitization so that people had information before the vaccine arrived.

> *Communities will need to be well sensitized. Chiefs, as community custodians and influential people, need to have the information. The ministry is now introducing other vaccines, but before giving out the vaccines, people need to be informed.* [FGD, health worker]

There was also demand for adults to be vaccinated, not just children:

> *We would love if the government would ensure that this vaccine reaches out to everyone, so we can stop this disease*. [FGD, female parent]

## Discussion

We found a broad spectrum of community views about the nature of influenza, including symptoms, types, severity, causes, prevention and treatment, and areas of both overlap and difference with biomedical understandings. Here we summarise these views and compare them to findings from other countries and biomedical understandings.

Participants often defined influenza as a condition characterized by nasal congestion or difficulty breathing, consistent with findings from South Africa [17]. Overlap in symptoms with other conditions meant some people saw influenza as a symptom or cause of conditions such as malaria and pneumonia. Influenza is an established cause of pneumonia (7,40), but not malaria. In addition, it was unclear whether participants just saw sneezing as an early symptom of pneumonia, or whether they understood that influenza infection increases a risk of developing pneumonia.

In relation to severity, some saw influenza as less dangerous than other conditions, because it was not fatal or did not disrupt work; similar to perceptions in Kenya (15), South Africa (17) and the UK (41). Others thought influenza was serious, feeling it did limit daily activities or could lead to death, especially when combined with other conditions. Young children and older people were perceived as at greater risk, consistent with medical understanding of these groups as more vulnerable to influenza complications (15,42).

In relation to causes of influenza, most people discussed dust as the cause, but hot and cold weather, poor hygiene and transmission between people were also mentioned. The association of dust or dirt with influenza was also found in Kenya and South Africa (15,17). Previous research suggests particulate matter contributes to respiratory infections in infants (43) and that dust could carry influenza viruses (44). However, participants tended to describe dust as making people sneeze, with only a few mentioning a link between germs and dust. This seems in part to reflect the broad interpretation of the Chichewa term chimfine, which as previously noted, goes beyond the biomedical definition of influenza to incorporate sneezing more generally. Beyond these immediate causes, some participants thought living conditions contributed to influenza, a perception supported by research in Malawi that indicates the significance of household crowding, poor sanitation and food insecurity (8).

Approaches to preventing influenza sometimes followed perceived causes, supporting the idea that health seeking reflects understandings of etiology (45). For example, ideas that contagion caused influenza were reflected in prevention strategies such as avoiding close contact with other people, a measure also reported in high-income settings (10,14). Similarly, keeping houses clean, not playing in dust and wearing warm clothes followed perceptions that influenza was caused by lack of hygiene, dust and cold, respectively. This supports the idea that how people manage illness reflects their understanding of its cause (45). However, respondents often held multiple and sometimes inconsistent beliefs about influenza. For example, people who identified dust as a cause sometimes also talked about catching influenza from other people, or mentioned warm clothes as a means of prevention, showing that explanatory illness models do not comprise only one fixed and coherent set of beliefs (46).

Treatment approaches included various home remedies such as hot water or herbs, with hospital visits made primarily when symptoms suggested potential malaria or pneumonia; similar findings are reported in Kenya (15). Several treatment strategies used by participants are consistent with biomedical guidance, including recommendations to relieve symptoms by drinking water and taking paracetamol [43,44]. The ideas of influenza as lacking treatment and of hospital visits as primarily required for more severe symptoms are also part of standard treatment guidance. Parents’ recognition that they should visit the hospital when there are signs of fever or rapid breathing perhaps reflects the effectiveness of health education in raising awareness about malaria and pneumonia in Malawi, something called for in previous studies [45,46].

Participants’ uncertainty about influenza and varied understanding of severity suggest that providing information for primary health workers and communities could support more confident influenza management. Health education could build on existing understanding. For example, while many participants described hygiene as a way to prevent influenza, this related primarily to household cleanliness, with only one participant mentioning hand washing and little attention to other steps to prevent spread of infection such as covering a sneeze. Beyond education, views from our participants and other research indicate the importance of addressing social determinants of health, such as overcrowded housing, within influenza control strategies [8,47,48]. Addressing these social determinants requires a multi-sectoral and long-term approach, the importance of which is increasingly recognised in international health discussions [42]. Our findings also suggest high acceptability of paediatric influenza vaccination, another potentially measure for influenza control (8,54). Parents’ enthusiasm for the influenza vaccine reflects Malawi’s high uptake of existing vaccinations (19).

The study had limitations. First, the broad definition of chimfine, encompassing influenza but also a runny nose, cold and sneezing more generally, complicates interpretation of the findings; parents may see dust as a cause of sneezing rather than of influenza. Research in English-speaking countries shows that the term ‘flu’ is used similarly, with the common cold or gastrointestinal symptoms sometimes described as flu (51,55). Further research could explore different interpretations of chimfine and community distinctions between the different conditions covered by this term. Second, the context of the influenza vaccine trial and a recognition that our study focused on influenza may have led parents to emphasize the severity of influenza. However, despite this context, many parents did describe influenza as less serious than other conditions. Third, we relied largely on reported practice rather than observation of actual influenza management. The critical incident narratives with parents who brought children to health centres provided concrete examples of treatment seeking, but did not include those who relied on home management. We sought to overcome this by asking all participants about recent experiences of influenza and how their treatment approach varies between illness episodes. Extended ethnographic observation within communities could provide further understanding. Finally, our findings are based on one district in Southern Malawi, and perceptions may be different in other socioeconomic, cultural or health system contexts.

## Conclusion

Our study provides the first detailed description of views about influenza in Malawi, and contributes to the literature on perceptions of influenza in low-income settings. Programmatically, the findings indicate the potential value of incorporating information about influenza within health education messages and providing information about influenza for community health workers. Our findings also suggest high acceptability of influenza vaccination for children, but acceptability of vaccination for other target groups including pregnant women and people with HIV requires further research. Finally, our research suggests the need for careful investigation of the broad meaning of chimfine and investigation of terminology used for influenza, to support clear communication between researchers, clinicians and communities.

## Acknowledgements

We would like to thank study participants for their contributions through participation and the FLUVAC study for permission to speak to their participants.

## References

1. Iuliano AD, Roguski KM, Chang HH, Muscatello DJ, Palekar R, Tempia S, et al. Estimates of global seasonal influenza-associated respiratory mortality: a modelling study. The Lancet [Internet]. 2017 Dec 13 [cited 2018 Mar 7];0(0). Available from: http://www.thelancet.com/journals/lancet/article/PIIS0140-6736(17)33293-2/abstract

2. Cohen C, Reubenson G. Influenza. 2017;1. Available from: http://www.springer.com/978-3-319-54032-0

3. McMorrow ML, Okitolonda Wemakoy E, Tshilobo JK, Emukule GO, Mott JA, Njuguna H, et al. Severe Acute Respiratory Illness Deaths in Sub-Saharan Africa and the Role of Influenza: A Case Series From 8 Countries. J Infect Dis. 2015 Sep 15;212(6):853–60.

4. Radin JM, Katz MA, Tempia S, Talla Nzussouo N, Davis R, Duque J, et al. Influenza Surveillance in 15 Countries in Africa, 2006–2010. J Infect Dis. 2012 Dec 15;206(suppl_1):S14–21.

5. Simoes EAF, Cherian T, Chow J, Shahid-Salles SA, Laxminarayan R, John TJ. Acute Respiratory Infections in Children. 2006 [cited 2017 May 30]; Available from: http://europepmc.org/abstract/med/21250360

6. Nyamusore J, Rukelibuga J, Mutagoma M, Muhire A, Kabanda A, Williams T, et al. The national burden of influenza-associated severe acute respiratory illness hospitalization in Rwanda, 2012-2014. Influenza Other Respir Viruses. 2017;

7. Cohen C, Simonsen L, Kang J-W, Miller M, McAnerney J, Blumberg L, et al. Elevated Influenza-Related Excess Mortality in South African Elderly Individuals, 1998–2005. Clin Infect Dis. 2010;51(12):1362–1369.

8. Ho A, Aston S, Jary H, Mitchell T, Alaerts M, Menyere M, et al. Impact of HIV on the burden and severity of influenza illness in adults in Malawi: a cohort and case-control study. The Lancet. 2016;387:S51.

9. Li J, Moore N, Akter S, Bleisten S, Ray P. mHealth for influenza pandemic surveillance in developing countries. In: System Sciences (HICSS), 2010 43rd Hawaii International Conference on [Internet]. IEEE; 2010 [cited 2017 May 29]. p. 1–9. Available from: http://ieeexplore.ieee.org/abstract/document/5428376/

10. Ferrante G, Baldissera S, Moghadam PF, Carrozzi G, Trinito MO, Salmaso S. Surveillance of perceptions, knowledge, attitudes and behaviors of the Italian adult population (18– 69 years) during the 2009–2010 A/H1N1 influenza pandemic. Eur J Epidemiol. 2011;26(3):211–219.

11. Liefooghe R, Baliddawa JB, Kipruto EM, Vermeire C, De Munynck AO. From their own perspective. A Kenyan community’s perception of tuberculosis. Trop Med Int Health. 1997;2(8):809–821.

12. Weiss MG. Cultural models of diarrheal illness: conceptual framework and review. Soc Sci Med. 1988;27(1):5–16.

13. Elledge BL, Brand M, Regens JL, Boatright DT. Implications of public understanding of avian influenza for fostering effective risk communication. Health Promot Pract. 2008;9(4_suppl):54S–59S.

14. Raude J, Setbon M. Lay perceptions of the pandemic influenza threat. Eur J Epidemiol. 2009;24(7):339–342.

15. Oria PA, Arunga G, Lebo E, Wong JM, Emukule G, Muthoka P, et al. Assessing parents’ knowledge and attitudes towards seasonal influenza vaccination of children before and after a seasonal influenza vaccination effectiveness study in low-income urban and rural Kenya, 2010–2011. BMC Public Health. 2013 Apr 25;13:391.

16. Rathi S, Gandhi H, Francis M. Knowledge and awareness about H1N1 flu in urban adult population of Vadodara, India. Electron Physician. 2011;3:392–95.

17. Wong KK, Cohen AL, Norris SA, Martinson NA, von Mollendorf C, Tempia S, et al. Knowledge, attitudes, and practices about influenza illness and vaccination: a cross‐sectional survey in two South African communities. Influenza Other Respir Viruses. 2016 Sep;10(5):421–8.

18. de Lataillade C, Auvergne S, Delannoy I. 2005 and 2006 seasonal influenza vaccination coverage rates in 10 countries in Africa, Asia Pacific, Europe, Latin America and the Middle East. J Public Health Policy. 2009;30(1):83–101.

19. PATH. PATH. Maternal Influenza Immunization: Perceptions of decision-makers, health care providers, and the community in Malawi. Seattle: PATH; 2016. 2016;

20. Health Sector Strategic Plan II 2017-2022 - Google Search [Internet]. [cited 2018 Mar 22]. Available from: https://www.google.com/search?q=Health+Sector+Strategic+Plan+II+2017-2022&ie=utf-8&oe=utf-8&client=firefox-b-ab

21. Schoub BD. Surveillance and management of influenza on the African continent. Expert Rev Respir Med. 2010;4(2):167–169.

22. Duque J, McMorrow ML, Cohen AL. Influenza vaccines and influenza antiviral drugs in Africa: are they available and do guidelines for their use exist? BMC Public Health [Internet]. 2014 Dec [cited 2018 May 28];14(1). Available from: http://bmcpublichealth.biomedcentral.com/articles/10.1186/1471-2458-14-41

23. Sambala EZ, Manderson L. Policy perspectives on post pandemic influenza vaccination in Ghana and Malawi. BMC Public Health. 2017 Feb 28;17:227.

24. Divala TH, Kalilani-Phiri L, Mawindo P, Nyirenda O, Kapito-Tembo A, Laufer MK. Incidence and Seasonality of Influenza-Like Illnesses Among Pregnant Women in Blantyre, Malawi. Am J Trop Med Hyg. 2016;95(4):915–917.

25. Ho A, Aston SJ, Jary H, Mitchell T, Alaerts M, Menyere M, et al. Impact of HIV on the burden and severity of influenza illness in Malawian adults: a prospective cohort and parallel case-control study. Clin Infect Dis. 2017;

26. National Statistical Office. MDG Endline Survey 2014: Chikwawa. Zomba, Malawi: National Statistical Office; 2016.

27. Government of Malawi. Integrated Household Survey 2010-2011. Zomba, Malawi: National Statistical Office; 2012.

28. Dzinjalamala F. Epidemiology of malaria in Malawi. Epidemiol Malawi. 2009;203:21.

29. Msyamboza K, Ngwira B, Banda R, Mkwanda S, Brabin B. Sentinel surveillance of Lymphatic filariasis, Schistosomiasis, Soil transmitted helminths and Malaria in rural southern Malawi. Malawi Med J [Internet]. 2010 [cited 2017 Aug 7];22(1). Available from: https://www.ajol.info/index.php/mmj/article/view/55901

30. Nyirenda TS, Molyneux ME, Kenefeck R, Walker LS, MacLennan CA, Heyderman RS, et al. T-Regulatory Cells and Inflammatory and Inhibitory Cytokines in Malawian Children Residing in an Area of High and an Area of Low Malaria Transmission During Acute Uncomplicated Malaria and in Convalescence. J Pediatr Infect Dis Soc Pp. 2015;1:10.

31. National Statistical Office. National Statistical Office (NSO) [Malawi] and ICF. 2017. Malawi Demographic and Health Survey 2015-16. Zomba, Malawi, and Rockville, Maryland, USA. NSO and ICF [Internet]. 2016 [cited 2017 Dec 7]. Available from: https://dhsprogram.com/pubs/pdf/FR319/FR319.pdf

32. Martindalea S, Mkwandab SZ, Smitha E, Molyneuxa D, Stantona MC, Kelly-Hopea LA. Quantifying the physical and socio-economic burden of filarial lymphoedema in Chikwawa District, Malawi. prospects. 2014;7(8):20–21.

33. Stanton M, Yamauchi M, Mkwanda SZ, Ndhlovu P, Matipula DE, Mackenzie C, et al. Measuring the physical and economic impact of filarial lymphoedema in Chikwawa district, Malawi: a case-control study. Infect Dis Poverty. 2017;6(1):28.

34. Kambauwa G, Gama S, Makumba W, Chaula K. Baseline Survey Report. 2010;

35. Peterson I. Influenza vaccine response in children 6-59 months residing in malaria endemic area of Malawi [Internet]. isrctn. 2016 [cited 2017 Aug 7]. Available from: http://www.isrctn.com/ISRCTN51364654

36. Norad. Local Perceptions, Participation and Accountability in Malawi’s Health Sector [Internet]. 2013 [cited 2018 Mar 22]. Available from: http://www.opml.co.uk/sites/default/files/Norad%20-%20Local%20Perceptions%2C%20Participation%20and%20Accountability%20in%20Malawi%27s%20Health%20Sector_0.pdf

37. Byrne M. Critical incident technique as a qualitative research method. AORN J. 2001 Oct 1;74(4):536–9.

38. Paas S. Chichewa-English/English-Chichewa Dictionary - Steven Paas - Oxford University Press [Internet]. 2013 [cited 2017 Oct 18]. Available from: https://global.oup.com/academic/product/chichewa-englishenglish-chichewa-dictionary-9780190416591?lang=en&cc=fr

39. Malawi Meteorological Services [Internet]. [cited 2018 Mar 22]. Available from: https://www.metmalawi.com/climate/climate.php

40. Brooks WA, Goswami D, Rahman M, Nahar K, Fry AM, Balish A, et al. Influenza is a major contributor to childhood pneumonia in a tropical developing country. Pediatr Infect Dis J. 2010;29(3):216–221.

41. Evans MR, Prout H, Prior L, Tapper-Jones LM, Butler CC. A qualitative study of lay beliefs about influenza immunisation in older people. Br J Gen Pr. 2007;57(538):352–358.

42. Ghebrehewet S, MacPherson P, Ho A. Influenza. BMJ. 2016 Dec 7;355:i6258.

43. Gurley ES, Homaira N, Salje H, Ram PK, Haque R, Petri W, et al. Indoor exposure to particulate matter and the incidence of acute lower respiratory infections among children: A birth cohort study in urban Bangladesh. Indoor Air. 2013;23(5):379–386.

44. Chen P-S, Tsai FT, Lin CK, Yang C-Y, Chan C-C, Young C-Y, et al. Ambient Influenza and Avian Influenza Virus during Dust Storm Days and Background Days. Environ Health Perspect. 2010;118(9):1211.

45. Green AR, Carrillo JE, Betancourt JR. Why the disease-based model of medicine fails our patients. West J Med. 2002;176(2):141.

46. Williams B, Healy D. Perceptions of illness causation among new referrals to a community mental health team:“explanatory model” or “exploratory map”? Soc Sci Med. 2001;53(4):465–476.

47. CDC. Wash your hands often and right way. [Internet]. Centers for Disease Control and Prevention. 2015 [cited 2017 Nov 2]. Available from: http://www.cdc.gov/flu/consumer/caring-for-someone.htm

48. NHSUK. Flu - NHSUK [Internet]. [cited 2017 Nov 2]. Available from: https://www.nhs.uk/conditions/flu/#what-to-do

49. Lutala PM, Mzumara S, Mlenga M, Talipu R, Kasagila E. Pneumonia in rural Malawians under five years old: Treatment outcomes and clinical predictors of death on admission. Afr J Prim Health Care Fam Med. 2009;1(1):1–6.

50. Masangwi SJ, Ferguson NS, Grimason AM, Morse TD, Kazembe LN. Care-seeking behaviour and implications for malaria control in southern Malawi. South Afr J Epidemiol Infect. 2010;25(4):22–26.

51. McCombie SC. Folk flu and viral syndrome: An epidemiological perspective. Soc Sci Med. 1987 Jan 1;25(9):987–93.

52. Quinn SC, Kumar S. Health inequalities and infectious disease epidemics: a challenge for global health security. Biosecurity Bioterrorism Biodefense Strategy Pract Sci. 2014;12(5):263–273.

53. CSDH. Closing the gap in a generation: health equity through action on the social determinants of health: Commission on Social Determinants of Health final report. World Health Organization; 2008.

54. CDC. Prevention and Control of Influenza [Internet]. 2003 [cited 2018 Mar 21]. Available from: https://www.cdc.gov/mmwr/preview/mmwrhtml/rr5208a1.htm

55. Jutel A, Banister E. “I was pretty sure I had the ‘flu”: Qualitative description of confirmed-influenza symptoms. Soc Sci Med. 2013 Dec 1;99(Supplement C):49–55.

